# AN OPTIMIZED METHOD TO DIFFERENTIATE HL60 CELLS INTO NEUTROPHIL-LIKE CELLS

**DOI:** 10.64898/2026.01.06.697988

**Authors:** Samuel P. Collie, Grant Yu, Suraj S. Rawat, Catharina Rypstra, Carole A. Parent

## Abstract

The HL60 promyelocytic leukemia cell line is widely used to investigate neutrophil biology due to its genetic tractability and accessibility. HL60 cells can be differentiated into neutrophil-like cells using all-trans retinoic acid (ATRA) or dimethyl sulfoxide (DMSO) treatment. However, these approaches produce cells that lack critical features of mature granulocytes, such as robust chemotaxis with ATRA treatment or multilobed nuclear morphology with DMSO treatment. To overcome these limitations, we developed a sequential differentiation protocol - ATRA for one day followed by DMSO for four days (A1D4) - which yields neutrophil-like cells that more faithfully recapitulate the morphology and functionality of mature human neutrophils. A1D4-differentiated HL60 cells display segmented nuclei with low lamin A/C expression and demonstrate strong chemotactic, oxidative burst, and phagocytic activity. This protocol combines the advantages of individual compounds, producing cells that closely mimic primary neutrophils in both phenotype and function.

**SUMMARY SENTENCE:** The authors provide an improved method to differentiate HL60 cells into neutrophil-like cells that faithfully recapitulates the morphology and functionality of mature human neutrophils.

## INTRODUCTION

Polymorphonuclear neutrophils represent the body’s first line of defense to injury and infection. They are the most abundant type of leukocytes, accounting for approximately 60% of the total leukocyte population in human blood^1^. Neutrophils circulate through the bloodstream until they sense chemical cues from injured/infected tissues, such as the formylated peptide N-formyl-Met-Leu-Phe (fMLF) which is released from bacteria and injured cells. In response, they transmigrate through the endothelial cell wall and chemotaxe to the affected area, where they perform critical functions including phagocytosis, degranulation, and the generation of reactive oxygen species (ROS) to eliminate pathogens and clear cellular debris^2^. Neutrophils also play essential roles in the pathophysiology of a broad range of diseases, including chronic inflammatory and autoimmune diseases, and cancer^3^. Popular models for studying neutrophils include neutrophils isolated from the blood of human donors, neutrophils isolated from the bone marrow of mice, induced pluripotent stem cells and chemically inducible cell lines^4^. Neutrophils from human blood are directly applicable to human health but are not easily accessible. In addition, these cells do not survive long *ex vivo* and are not amenable to genetic manipulations. Murine neutrophils are more readily available and, with the easy access to genetically modified mice, represent a popular model to dissect their function. However, murine neutrophils exhibit significant differences from their human counterparts. For example, mice have a lower proportion of neutrophils in their blood (<30% of leukocytes) compared to humans and they differ in their signaling pathways, granule protein composition and nuclear morphology^5,6^. Furthermore, the limited number of cells isolated from the bone marrow limit the ability to perform biochemical analyses. Finally, induced pluripotent stem cells offer a genetically amenable and phenotypically relevant alternative. However, protocols for producing mature neutrophils from stem cells are expensive and lengthy – making accessibility a significant drawback^7^. In contrast, human cell lines offer a valuable alternative, as they are genetically tractable, physiologically relevant, and readily available^8,9^.

Among cell line models, the most well established is the human leukemia line, HL60^4,10^. HL60 cells were isolated from a patient suffering from acute myeloblastic leukemia with maturation^10,11^. Another popular human leukemic cell line, PLB-985, was shown to be a subclone of HL60^12^, underscoring the prevalence of the HL60 cell line in neutrophil research. Importantly, HL60 cells can be readily differentiated into granulocytic lineages to reproducibly produce >90% differentiated cells^13^. However, these cells do not fully recapitulate the morphology and function of mature neutrophils isolated from human blood^14^. Two of the most widely used differentiation agents for inducing differentiation are the polar organic solvent, dimethyl sulfoxide (DMSO)^15^ and the vitamin A derivative, all-trans retinoic acid (ATRA)^16,4^. Although both DMSO and ATRA treatment for five to six days lead to morphological and protein expression changes associated with mature granulocytes, these factors induce slightly different phenotypic states. DMSO differentiated cells do not adopt a lobulated nuclear morphology, whereas ATRA differentiated cells express low levels of chemoattractant receptors^13,17^ and exhibit poor chemotactic responses^4^. In general, ATRA-differentiated cells have a more mature granulocytic phenotype transcriptionally, whereas DMSO-differentiated cells have increased immune responses^18^.

By combining ATRA and DMSO treatment, both the mature phenotypic state of ATRA-differentiated cells and the robust immune response observed with DMSO-differentiated cells could be retained. ATRA is known to promote differentiation initially through transcriptional changes mediated by its binding to the nuclear Retinoic Acid Receptor alpha (RARα)^19^. DMSO mediates differentiation through a combination of changes in membrane fluidity^20^, chromatin remodeling^21^, and upregulation of Ras and Protein Kinase C (PKC) signaling pathways^22,23^. The combined use of DMSO and ATRA to optimize HL60 differentiation has been investigated, but its effects on cellular morphology and function has not been fully characterized^24–26^. In this study, we developed a sequential differentiation protocol in which ATRA treatment is followed by DMSO exposure, that gives rise to neutrophil-like cells exhibiting a lobulated nuclear morphology and functional chemotaxis, phagocytosis, and oxidative burst capacity. This optimized approach provides an easily accessible and physiologically relevant cell line model of neutrophils, enabling detailed studies of their fundamental biology.

## METHODS

### Cell Culture

HL60 cells were purchased from ATCC (CCL-240) and were cultured in Iscove’s Modified Dulbecco’s Medium IMDM (Gibco-12440-053) with 10% heat-inactivated fetal bovine serum (HI-FBS) and a penicillin–streptomycin antibiotic cocktail. Cells were cultured in 5% CO2 humidified air at 37°C and tested to exclude mycoplasma contamination. Cells were passaged every 2–3 days and cells from no more than 12 passages were used for all experiments.

### Isolation of neutrophils from human blood

Neutrophils were isolated from the peripheral blood of healthy, human donors using a protocol adapted from Kremserova and Nauseef^27^. Peripheral blood was obtained through the Platelet, Pharmacology and Physiology Core facility at the University of Michigan, which recruits and collects blood of anonymous healthy donors by venipuncture. This protocol was approved by institutional review board (HUM00107120) to supply de-identified blood samples for research purposes. Therefore, we do not have access to any identifiable donor information under the coverage of the Health Insurance Portability and Accountability Act. Participants consented to donate their blood as a biological sample for research purposes and were financially compensated. Donors were additionally required to verify that they did not take aspirin for 7 days and NSAIDS for 48h before venipuncture.

### HL60 Differentiation with ATRA and DMSO

HL60 cells were differentiated with 1.3% DMSO in 10% HI-FBS containing IMDM media for 6 days as previously described^8^ or alternatively with 1µM or 250nM ATRA in 10% HI-FBS containing IMDM media for 6 days^24^. Additionally for our new method, HL60 cells were differentiated using 250nM ATRA in 5% HI-FBS containing IMDM media with 6.7ng/mL Selenium, 10µg/mL Insulin, and 5.5µg/mL Transferrin (ITS) at a concentration of 0.4 million cell/mL. After 24 h, cells were centrifuged and re-plated with 1.3% DMSO in 5% HI-FBS IMDM with ITS for two days. Then, cells were centrifuged and re-plated in double the volume of fresh 1.3% DMSO containing IMDM media with 5% HI-FBS IMDM and ITS for two more days. On the final day of differentiation, differentiated HL60 (dHL60) cells were counted and assessed for viability using Trypan blue before beginning experiments. For all experiments Hank’s Balanced Salt Solution with calcium, magnesium, and 25mM D-Glucose was used to resuspend cells – hereafter referred to as mHBSS.

### High content confocal microscopy of activated dHL60 cells

24-well cell culture plates were coated with 25µg/mL fibrinogen (Sigma-Aldrich F3879) for at least 1 h at 37°C. 1.5 x 10^6^ dHL60s or human neutrophils were plated in each well in HBSS (Gibco 14025092) and allowed to settle for 10 min at 37°C. Cells were uniformly stimulated with LTB_4_ (Cayman Chemical 20110; 100 nM) for 15 min at 37°C, fixed with 4% PFA (Electron Microscopy Sciences 15714) in DPBS for 15 min at room temperature, blocked with DPBS containing 2% goat serum, 0.2% saponin (Sigma-Aldrich 84510), and incubated with antibodies against FLAP (ThermoFisher MA5-37933, 1:300), LBR (Abcam ab32535, 1:300), 5-LO (Abcam ab169755, 1:200), or LMNA (Cell Signaling Technologies 4777S, 1:100) at 4°C for overnight. Samples were washed with DPBS for 5 min 3X before adding DAPI (10µg/µL), Goat anti-rabbit AlexaFluor488 and Goat anti-mouse AlexaFluor568 (Invitrogen A11008/A11031, 1:500) for 1 h at room temperature. After washing in DPBS for 5 min 3 times, samples were imaged using the Yokogawa CellVoyager (CV8000), a high content confocal microscope, with a 40x water-immersion objective lens and maximum-intensity projection processing. Images were analyzed using an in-house CellProfiler pipeline to identify and segment nuclei objects, measure fluorescent intensities in the nucleus, and quantify nuclear morphology.

### Underagarose Migration Assay

The migration assay was performed as previously described^8^. Briefly, glass bottom dishes were coated with 1% BSA for at least 1 h at 37°C and then cast with a 0.5% agarose in DPBS:HBSS (1:1). The gel was allowed to solidify for 15 min at room temperature then 45 min at 4°C. A 3x3 array of 1 mm diameter wells 2 mm apart from each other was punched into the gel. 50,000 Hoechst-stained cells suspended in 5 µL of mHBSS was added to each well in the left and right columns and allowed to settle for 10 min at 37°C. 100 nM fMLF was added to the middle column, creating a gradient of 50 pM µm^-1^, as previously described^28^, and displayed in Figure 1B. Chemotaxis was recorded using the Zeiss Axio Observer.Z1 with a 10x objective lens for 90 min at 37°C. Migrating cells were tracked using the TrackMate plugin in ImageJ and tracks were analyzed using an in-house MatLab script as previously described^29^.

**Figure 1.**
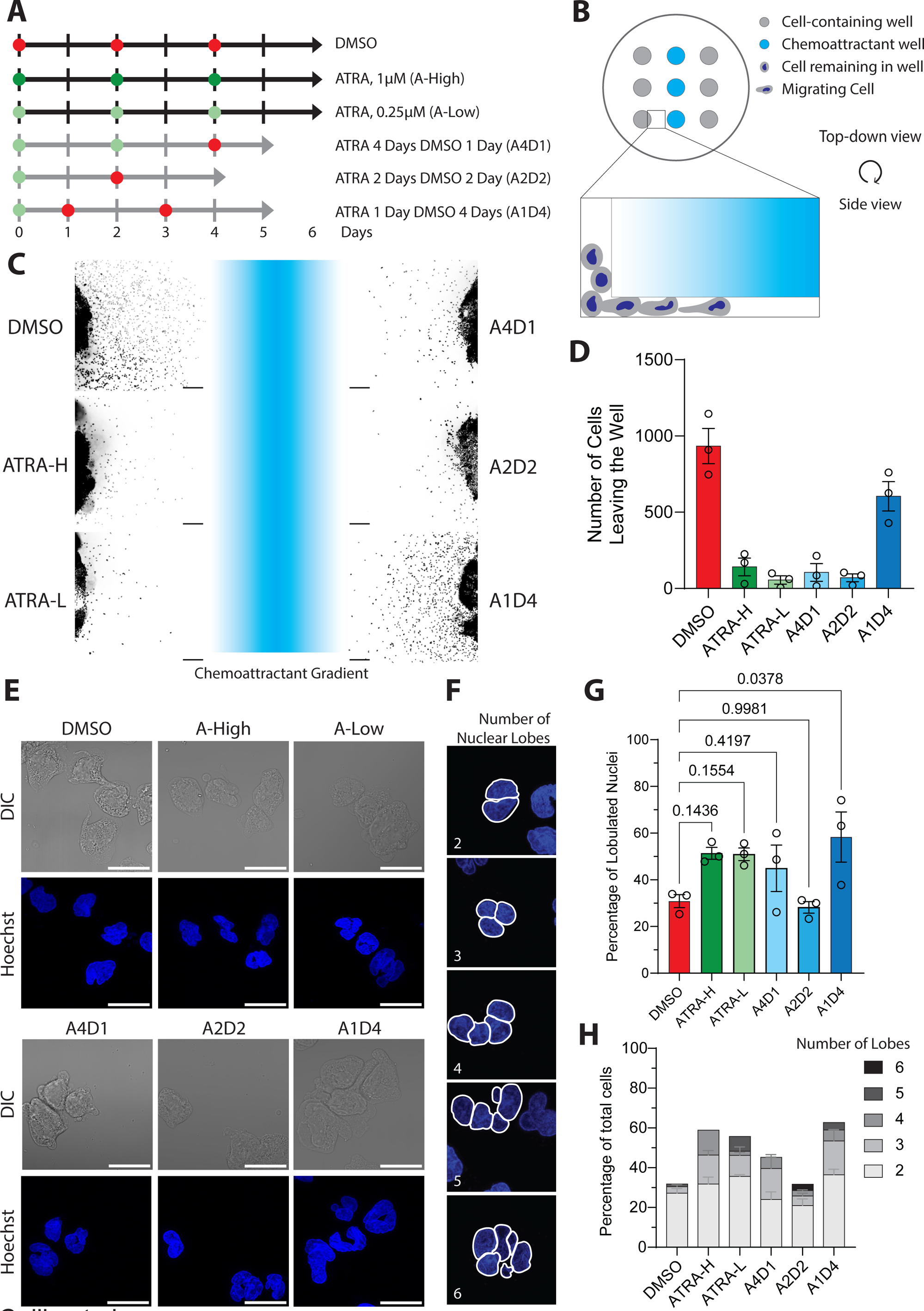
HL60 differentiation with ATRA and DMSO for migration and nuclear lobulation. **A.** Schematic of the treatment conditions tested to optimize HL60 differentiation. Differentiation proceeds over a period of 4-6 days depending on condition. Black bars indicate HL60 cells cultured in 10% HI-FBS, grey bars indicate cells cultured in 5% HI-FBS with supplemented ITS. Red circles are treatment with 1.3% DMSO, dark green circles are treatment with 1µM ATRA, and light green circles are treatment with 250nM ATRA. **B.** Cartoon depicting the underagarose migration assay. Cells (grey) migrate underneath agarose towards a chemical gradient (light blue). **C.** Representative widefield microscopy images of dHL60 cells that migrated for 60 min toward fMLF, fixed and stained with Hoechst. Visual representation of the chemoattractant gradient is shown in between the cells in light blue. Scale bar is 200µm. **D.** Bar graph showing the number of dHL60 cells that migrated from the well toward the fMLF gradient. Data are presented as mean ± s.e.m. N=3. **E.** Airyscan microscopy images of fixed dHL60 cells that migrated for 60 min toward fMLF, fixed and stained with Hoechst. **F.** Representative airyscan confocal images of dHL60 cells with outlines of manually counted nuclear lobes. **G.** Bar graph showing the percentage of lobulated nuclei (more than 1 nuclear lobe) of the total cells imaged, at least 75 cells for each treatment. Bar represents the mean ± s.e.m. *P* values calculated using repeated measures (RM) one-way ANOVA N=3. **H.** Stacked bar graph showing the distribution of lobulation amongst cells with lobulated nuclei. Data are presented as mean ± s.e.m.

### Flow Cytometry

#### For Differentiation Markers

1 x 10^6^ dHL60 cells were washed with 500µL HBSS and resuspended in 0.5mL of FACS buffer (HBSS +2% FBS + 1mM EGTA, 0.05% Sodium Azide) in BSA-coated tubes. Cells were incubated with mouse anti-CD11b antibody conjugated to Allophycocyanin (APC) (1:5) or an isotype control and stimulated with either 1µM fMLF-AlexaFluor488 or 1µM unconjugated fMLF on ice in the dark for 30 min. This will allow for the readout of the expression of CD11b and the receptor for fMLF – FPR1 – on the cell surface which are both associated with mature neutrophils^30,31^. Cells were then centrifuged at 300 x g for 3 min at 4°C, washed with 100µL ice cold FACS buffer 3 times and fixed in 300µL of 4% PFA in HBSS for 10 min at room temperature. Fixed cells were washed and resuspended in HBSS before acquisition.

#### For Reactive Oxygen Species

0.3 × 10⁶ cells were resuspended in 1 mL HBSS containing 50 mM glucose, and 2.5 µM DCFDH, a chemiluminescent redox probe^32^, in BSA-coated tubes. Cells were incubated for 15 min at 37 °C under gentle rotation (10 rpm), stimulated with 100nM fMLF for 2 min and centrifuged at 500 × g for 5 min at 4 °C. The supernatant was discarded, cells were washed twice with ice-cold 1× DPBS, the pellet was resuspended in 400 µL ice-cold 1× DPBS and analyzed immediately by flow.

#### For Apoptosis Assay

0.3 × 10⁶ cells were seeded in 1 mL apoptosis binding buffer containing 1 µg/mL propidium iodide and 3 µL/mL ApoScreen® Annexin V–FITC in BSA-coated tubes. This combination allows for a readout of phosphatidylserine on the cell surface through Annexin V and the permeability of the cell membrane using propidium idodide. Taken together this can be used to detect apoptotic state of cells at a population level^33^. Cells were incubated on ice for 15 min in the dark with intermittent inversion and centrifuged at 500 × g for 5 min at 4 °C. The supernatant was discarded, cells were washed twice with ice-cold DPBS, the pellet was resuspended in 400 µL ice-cold 1× DPBS, transferred to flow cytometry tubes, and analyzed immediately. Data was acquired on either LSR Fortessa, Sony ID7000, Sony SH800 or Bio-Rad Ze#5 analyzers, recording 100,000 events per sample.

### Phagocytosis Assay

To model phagocytic capacity, internalization of fluorescent zymosan was assesed^34^. 2×10⁶ zymosan bio-particles conjugated to AlexaFluor 594 (Invitrogen Z23374) were opsonized in 1 mL of opsonization medium (10% FBS in 1X HBSS with 140mg/L CaCl_2_ and 100mg/L MgCl_2_) for 30 min at 37 °C under gentle rotation (10 rpm). The suspension was centrifuged at 4,000 × g for 5 min and the pellet was washed once with prewarmed 1X DPBS before resuspension in 50 µL HBSS supplemented with 50 mM glucose. In parallel, 8-well chamber slides were coated with 25 µg mL⁻¹ fibrinogen and 10 µg mL⁻¹ fibronectin for 1 h at 37 °C. The coating solution was removed, 0.2 × 10⁶ cells were seeded in 200 µL HBSS + Ca²⁺ + 50 mM glucose, adhesion was allowed for 10–15 min, and the chambers were placed on ice. Opsonized particles were then added to cells placed on ice, distributed evenly for 5 min, and the chambers were incubated at 37 °C for 15 min. Non-adherent cells and particles were removed by gentle washing 3X with warm DPBS. Cells were fixed with 4% paraformaldehyde in prewarmed 1× PHEM buffer for 20 min at room temperature, washed once with DPBS, and stained with phalloidin-488 (1 U mL⁻¹) and Hoechst for 15 min at room temperature in the dark. Cells were washed 3X with DPBS before mounting with mounting medium for imaging using a Zeiss LSM 880 microscope. Images were analyzed using a Cell Profiler pipeline that segmented cells based on Hoechst and phalloidin staining and the number of zymosan bioparticles internalized in each cell was recorded.

## RESULTS

### Sequential treatment of ATRA and DMSO produces differentiated HL60 cells with lobulated nuclei and robust migration towards fMLF

To optimize the differentiation of HL60 cells into neutrophil-like cells, we sequentially treated cells with ATRA and DMSO for different times and compared this to cells treated with each of the agents alone (Fig. 1A). 1.3% DMSO (D) and 1µM ATRA (ATRA-H) for 6 days conditions represent standard conditions for the differentiation of HL60 cells using DMSO and ATRA^4^. The 250nM ATRA (ATRA-L) was included as it is closer to the critical concentration needed to induce differentiation^35^. These conditions, in IMDM media supplemented with 10% fetal bovine serum (Fig. 1A, black lines), were compared to conditions where HL60 cells were differentiated in reduced (5%) serum conditions supplemented with insulin, transferrin and selenium (ITS) as follows: (i) 4 days with 250nM ATRA then 1 day of DMSO (A4D1), (ii) 2 days with 250nM ATRA then 2 days of DMSO (A2D2), or (iii) 1 day with 250nM ATRA then 4 days of DMSO (A1D4) (Fig. 1A, gray lines). Reduced serum and ITS supplementation was used in line with previous studies showing increased HL60 cell survival during differentiation and reduced background activation from serum-related factors^36,37^. Since nuclear morphology and chemotactic response represent hallmarks of neutrophil function, we used these two characteristics to compare the six differentiation conditions. We tested chemotactic ability using an under agarose migration assay where cells migrate underneath a bed of agarose towards a gradient of chemoattractant (Fig. 1B)^8^. We utilized the formyl peptide fMLF as a strong neutrophil chemoattractant and assessed migration using live cell microscopy (Fig. 1C and Supplementary Video 1). We found that only the DMSO and A1D4 differentiated HL60 cells (dHL60) exhibit chemotactic activity, showing a substantial number of cells migrating towards the fMLF signal (Fig. 1D). This is in line with previous studies showing that ATRA-H differentiated cells do not express chemoattractant receptors or migrate well compared to DMSO differentiated cells^13,38^. However, limiting ATRA treatment to one day, followed by 4 days of DMSO appears to alleviate this problem as we found that A1D4 treated cells show robust chemotactic activity (Fig. 1D). We next measured the nuclear morphology of the dHL60 cells migrating under agarose using Hoechst-stained nuclei as a guide (Fig. 1E). Nuclear lobes were counted manually after blinding (Fig. 1F) and we found that ATRA-H and ATRA-L dHL60 cells trended towards exhibiting more multilobulated nuclei, compared to DMSO dHL60 cells, as previously reported^39^. However, we found that treatment with ATRA for one day was enough to induce nuclei lobulation: A1D4 dHL60 cells had a significantly higher percentage of lobulated nuclei (Fig. 1G) as well as more cells with three or more lobes (Fig. 1H), compared to DMSO dHL60 cells.

### A1D4 and DMSO dHL60 cells have similar viability and granulocyte surface marker expression

It has been reported that ATRA-H dHL60 cells have a reduced proliferative capacity compared to DMSO dHL60 cells and are highly susceptible to cell death by apoptosis^40,41^. To investigate cell viability, we used flow cytometry and measured propidium iodide and AnnexinV staining in HL60 cells differentiated with either DMSO, ATRA-H or A1D4 methods (Fig. 2A). Consistent with previous reports, we found that ATRA-H differentiated cells have significantly reduced viability compared to DMSO differentiated cells. Yet, the A1D4 treated cells showed comparable viability to DMSO treated cells (Fig. 2B-D). This finding indicates that reducing the concentration and duration of ATRA treatment helps to alleviate its toxic effects on HL60 cells. Since A1D4 cells are comparable to DMSO in terms of migration and viability but with increased nuclear lobulation, we chose to compare A1D4 dHL60 cells to DMSO dHL60 cells for the remainder of the experiments in this study.

**Figure 2.**
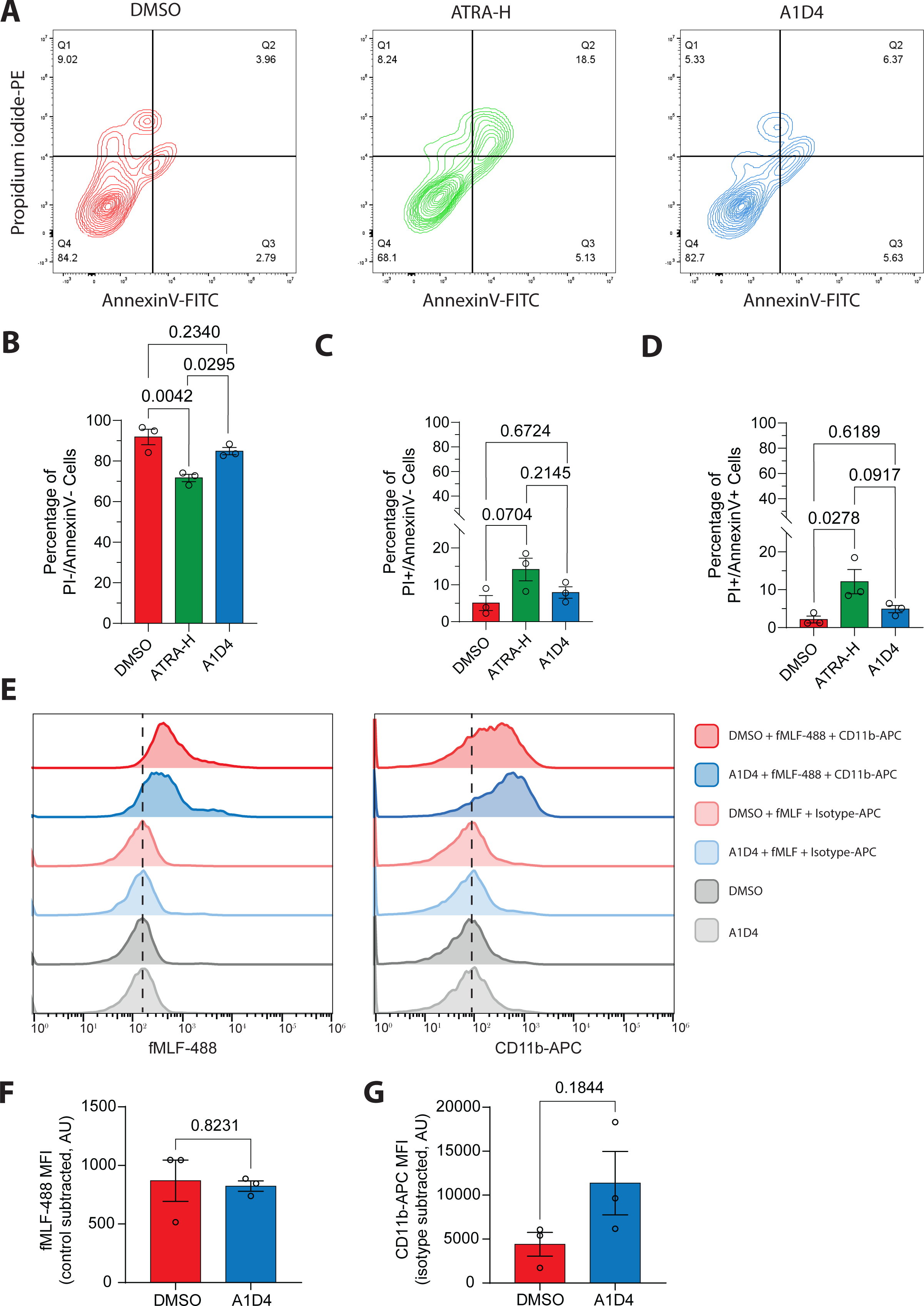
Viability and granulocytic maturity of A1D4 dHL60 cells. **A.** Representative contour diagrams showing propidium iodide-PE and AnnexinV-FITC fluorescence intensity of DMSO, ATRA-H, and A1D4 dHL60 cells. Values in the corner of each quadrant represent percentage of the population in that quadrant. **B-D.** Bar graphs showing percentage of cells PI-/AnnexinV-, PI+/AnnexinV-, and PI+/AnnexinV+ cells. Bar represents the mean ± s.e.m. *P* values calculated using RM one-way ANOVA. N=3. **E.** Representative histogram showing the intensity of fMLF-AlexaFluor488 (fMLF-488) and CD11b-APC in DMSO and A1D4 dHL60 cells. Non-fluorescent fMLF, isotype control-APC stained cells and unstained cells serve as negative controls. Dotted black line represents the peak of intensity in the negative controls. **F-G.** Bar graphs showing the fMLF-488 and CD11b-APC mean fluorescence intensity (MFI) with non-fluorescent fMLF and isotype control values subtracted. *P* values calculated using two-tailed Welch-corrected t test. Data are presented as mean ± s.e.m. N=3.

We next assessed the expression of cell surface markers observed with mature, fully differentiated granulocytes. For this we used flow cytometry targeting CD11b expression using a fluorescently tagged antibody, and formyl peptide receptor (FPR1) expression using the fluorescently tagged ligand fMLF-AlexaFluor488 (Fig. 2E). We found that A1D4 dHL60 cells express comparable levels of CD11b and FPR1 to DMSO dHL60 cells (Fig. 2F), with A1D4 cells showing a trend toward increased CD11b signal compared to DMSO dHL60 cells (Fig. 2G). These findings show that A1D4 dHL60 cells exhibit similar viability and expression of maturation markers compared to DMSO dHL60 cells.

### A1D4 dHL60 cells have higher phagocytotic capacity compared to DMSO dHL60 cells

Previous studies assessing the phagocytotic activity of dHL60 cells showed that ATRA-H dHL60 cells have a higher activity compared to DMSO dHL60 cells^25,39^. To assay phagocytic capacity, we measured uptake of opsonized zymosan bioparticles^34^ after 15 min of incubation using fixed cell imaging (Fig. 3A). We found that A1D4 dHL60 cells have significantly increased phagocytic activity (Fig. 3B) and an increase in the number of phagocytosed particles, compared to DMSO dHL60 cells (Fig. 3C). We next utilized the fluorogenic live cell ROS sensor, 2’,7’-dichlorodihydrofluorescein diacetate (DCFDA), which is oxidized to 2’, 7’-dichlorofluorescein (DCF) by ROS, to measure oxidative burst capacity in dHL60 cells by flow cytometry^32^. We found that A1D4 and DMSO dHL60 cells stimulated with fMLF for 2 min exhibit similar ROS generation (Fig. 3D-F). These results are in line with previous studies that reported no difference in the oxidative burst capacity of DMSO and ATRA-H differentiated HL60 cells^42,43^.

**Figure 3.**
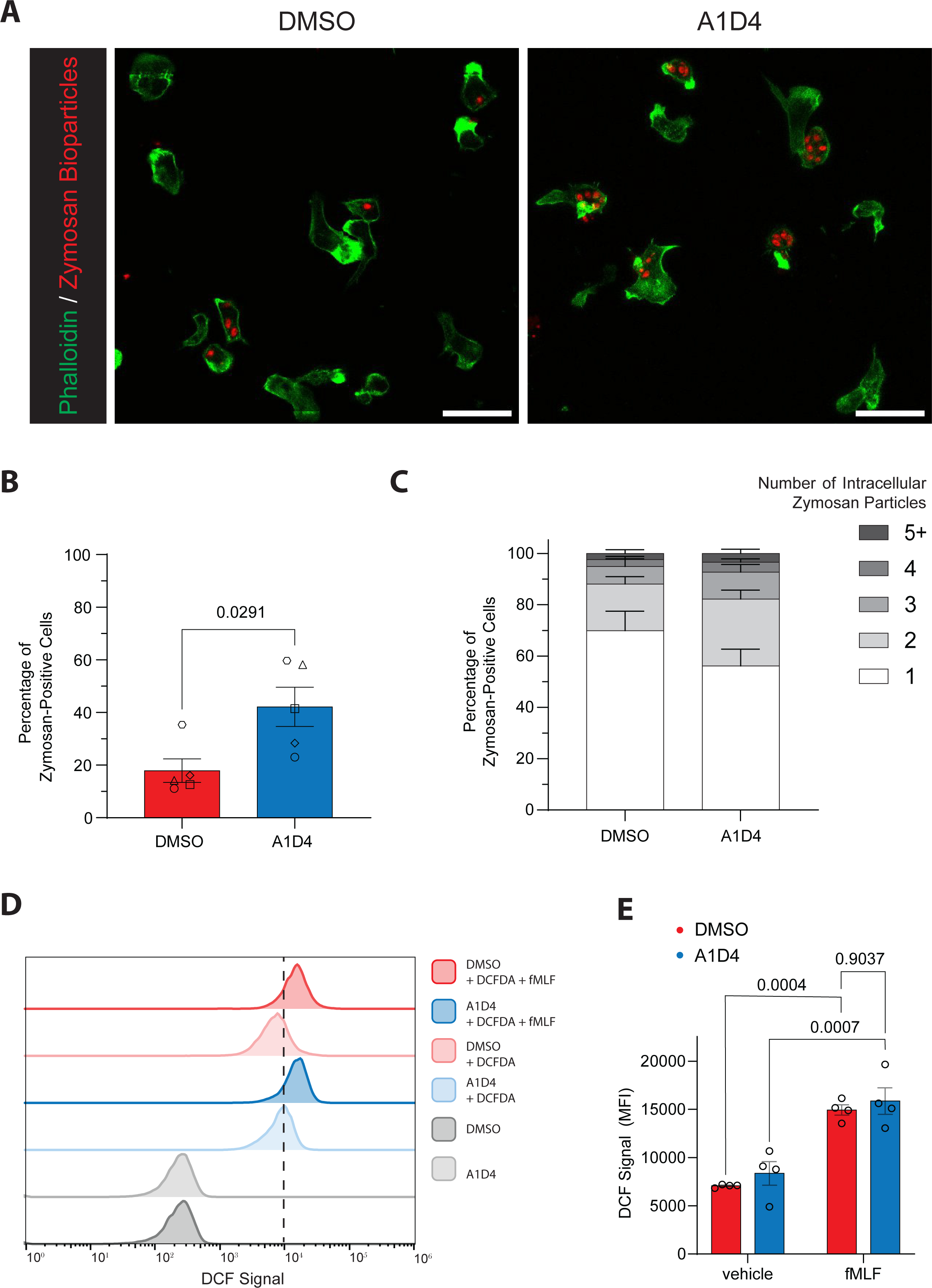
Phagocytosis and redox activity of activated A1D4 dHL60 cells. **A.** Representative confocal microscopy images of DMSO and A1D4 dHL60 cells visualized with phalloidin-AlexaFluor488 (green) and incubated with fluorescent zymosan bioparticles (red). Scale bar is 25µm. **B.** Bar graph showing the percentage of cells with an internalized zymosan bioparticle of total cells imaged. Data presented as mean ± s.e.m. *P* values calculated using two-tailed Welch-corrected t test. N=5. **C.** Stacked bar graph showing the number of internalized zymosan bioparticles amongst cells with internalized zymosan bioparticles. Data are presented as the mean ± s.e.m. **D.** Representative histogram showing the intensity of DCF in DMSO and A1D4 dHL60 cells stimulated with fMLF for 2 min. Unstimulated vehicle cells with DCFDA and unstained cells serve as negative controls. **E.** Bar graph showing the mean fluorescence intensity (MFI) of DCF in DMSO (red) and A1D4 (blue) dHL60 cells stimulated with fMLF or vehicle control. Data are presented as mean ± s.e.m. *P* values calculated using repeated measures (RM) one-way ANOVA. N=4.

### A1D4 dHL60 cells exhibit robust chemotactic responses

We next sought to further characterize the chemotactic ability of A1D4 dHL60 cells compared to DMSO dHL60 cells. To do so, we used live cell imaging to record dHL60 migration under agarose in response to gradients of the formyl peptide fMLF, the bioactive lipid leukotriene B_4_ (LTB_4_), and of the cytokine interleukin-8 (IL-8) (Supplementary Video 2). The migrating cells were tracked using the ImageJ plugin Trackmate^44^, analysis was performed using an in-house MatLab script, and chemotactic parameters including cell speed, Euclidean distance traveled and directionality were reported as previously described^29^ (Fig. 4A). In addition, we calculated the chemotactic fraction by dividing the number of cells migrating towards the chemoattractant by the total number of cells migrating toward and away from the chemoattractant well (Fig. 4B). The number of cells migrating towards fMLF and LTB_4_ was generally higher for both conditions compared to IL-8 as exemplified by the number of data points when each cell is plotted (Supplementary Fig. 1). It has been previously reported that in contrast to primary neutrophils, DMSO dHL60 cells express relatively low levels of the receptor for IL-8, CXCR1, when compared to the fMLF receptor, FPR1, and the LTB_4_ receptor, BLT1^37^. We found no significant differences in the chemotactic fraction, average cell speed, Euclidean distance traveled, and directionality between A1D4 and DMSO dHL60 cells migrating towards fMLF (Fig. 4C-F) or LTB_4_ (Fig. 4G-H). However, while A1D4 and DMSO cells responded equally to IL-8 (Fig. 4K-M), we did measure a reduction in directionality of A1D4 dHL60 cells toward IL-8, compared to DMSO dHL60 cells (Fig. 4N). Taken together, these results show that A1D4 dHL60 cells have similar chemotactic ability to DMSO dHL60 cells.

**Figure 4.**
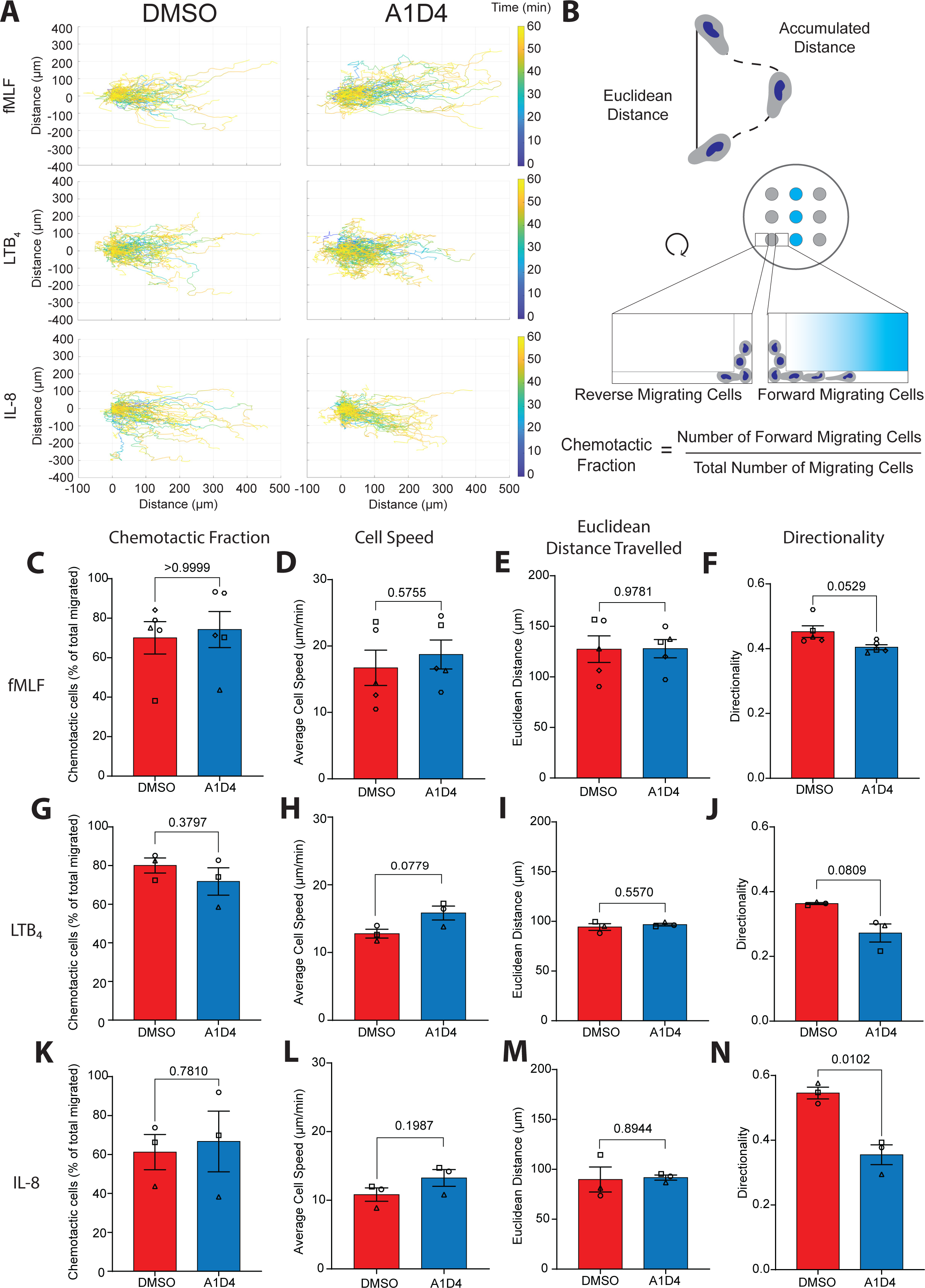
A1D4 dHL60 cells exhibit robust chemotactic responses. **A.** Graphs displaying tracks of 100 randomly selected DMSO or A1D4 dHL60 cells migrating towards fMLF, LTB_4_, or IL-8. Color-coded time scale is shown on the right. **B.** Cartoon depicting the accumulated distance and Euclidean distance travelled for a cell (top) and the underagarose assay and how chemotactic fraction is calculated (bottom). **C-N.** Bar graphs showing chemotactic parameters acquired from cells migrating for 60 min towards fMLF (**c-f**) N=5, LTB_4_ (**g-j**) N=3, IL-8 (**k-n**) N=3. For each chemoattractant the following parameters are displayed: chemotactic fraction (**c, g, k**), which indicates the proportion of cells migrating toward the chemoattractant gradient as described in **b;** average cell speed (**d, h, l**), which represents the means of all cells average speed during migration from each independent experiment; Euclidean distance travelled (**e, i, m**), which shows the means of all cells distance traveled in a straight line averaged for all cells across each independent experiment; and directionality (**f, j, n**), which represents the ratio of Euclidean distance to total accumulated distance traveled averaged for all cells across each independent experiment. *P* values calculated using two-tailed Welch-corrected t test. Data are presented as mean ± s.e.m.

### A1D4 dHL60 cells exhibit lobulated nuclei and lower lamin A/C expression

Nuclear segmentation and changes in the composition of nucleoskeletal proteins are hallmarks of mature granulocytes^45^. In particular, mature human neutrophils show a reduction in the expression of lamin A/C (LMNA/C), leaving B-type lamins (LMNB) as the dominant intermediate filament present in their nuclear lamina, and an increase in the presence of the inner nuclear transmembrane protein lamin B receptor (LBR)^46^. We and others have reported that DMSO dHL60 cells express LMNA/C along with LMNB1 and LMNB2^29,37^. In contrast, ATRA dHL60 cells have a lamin expression profile that is similar to human primary neutrophils, i.e. low levels of LMNA/C and high levels of LMNB1 and LMNB2^45,47^. Since we observed that A1D4 dHL60 cells exhibit multilobulated, segmented nuclei (Fig. 1E), we assessed the expression of nuclear proteins in these cells using Western blot analysis. We found that while A1D4 and DMSO dHL60 cells express similar levels of LBR, A1D4 differentiation leads to a consistent but non-significant reduction in the expression of both LMNA/C isoforms, compared to the DMSO differentiation method (Supplementary Fig. 2A-E). To assess nuclear shape and protein expression at the level of individual nuclei, we used high-throughput, confocal immunofluorescence microscopy on LTB_4_-uniformly stimulated A1D4 or DMSO dHL60 cells (Fig. 5A&B, Supplementary Fig. 1F&G). Using the CellProfiler software^48^, we computationally segmented nuclear objects from thousands of cells per condition and compared nuclear shape as well as nuclear protein expression by immunofluorescent signal. Using eccentricity and form factor measurements - geometric quantifications of shape irregularity that can be used to describe segmented nuclei (Fig. 5C)^49^ - we measured significantly more segmented nuclei in A1D4 compared to DMSO dHL60 cells (Fig. 5D&E). This is in line with our results from manually counting nuclear segmentation in migrating cells (Fig. 1E). Using immunofluorescence, we next determined the expression of LBR and LMNA/C in the nuclei of A1D4 or DMSO dHL60 cells. As observed by western blot analysis, we found no difference in LBR expression between A1D4 and DMSO dHL60 nuclei (Fig. 5F&I). However, we measured a significant decrease in LMNA/C expression in A1D4 dHL60 nuclei, compared with DMSO dHL60 nuclei (Fig. 5G&J). We also investigated the nuclear expression of Five-Lipoxygenase Activating Protein (FLAP), a endoplasmic reticulum and nuclear envelope transmembrane protein^29,50,51^ essential for the synthesis of LTB_4_ and the neutrophil swarming response^52^. We found that FLAP expression is significantly higher in A1D4 dHL60 cells compared to DMSO dHL60 cells (Fig. 5H&K). Together, these findings show that compared to DMSO, A1D4 treatment induces a more segmented nuclear morphology in dHL60 cells and an expression profile of nucleoskeletal proteins closer to human primary neutrophils.

**Figure 5.**
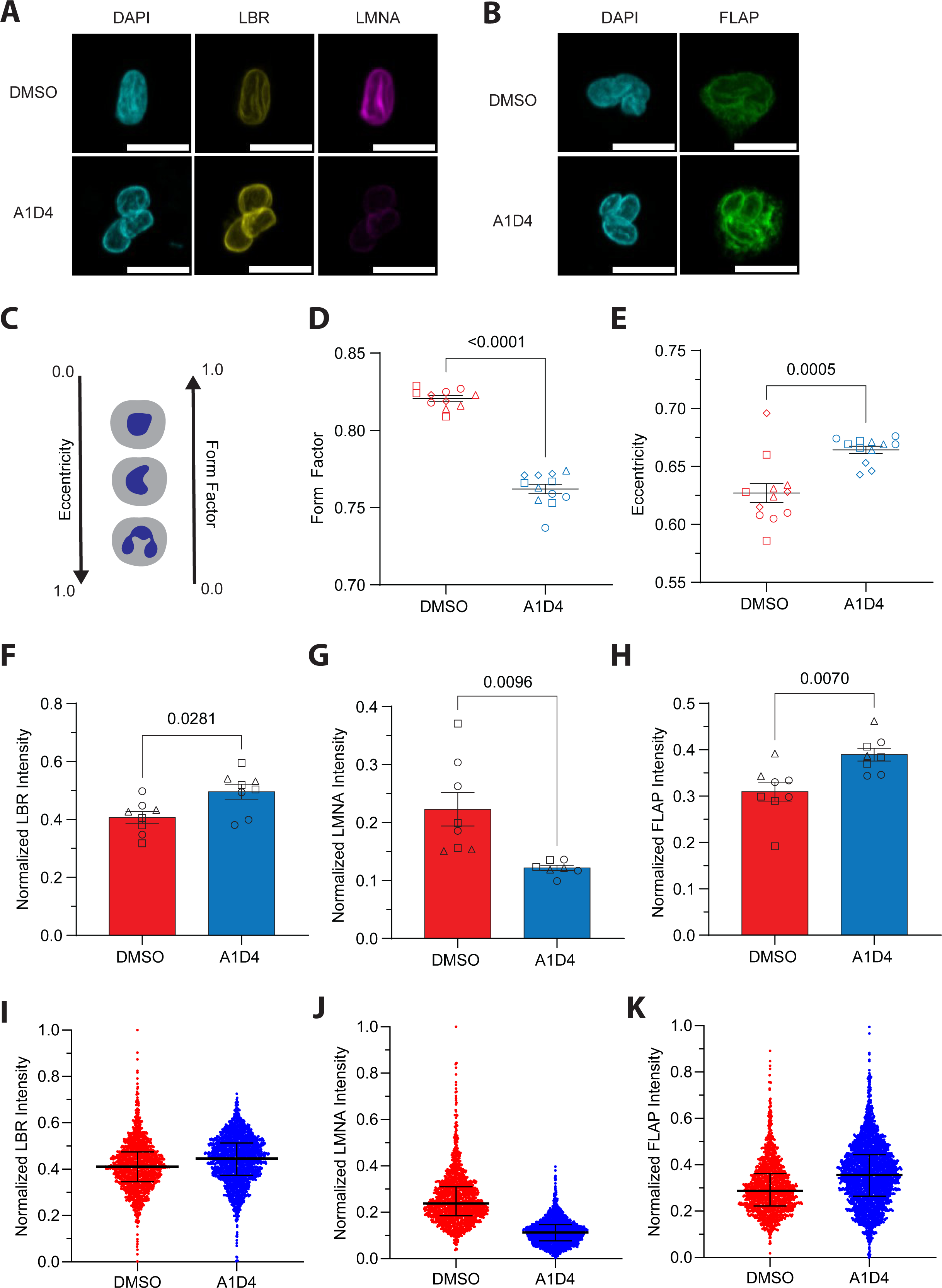
A1D4 dHL60 cells exhibit lobulated nuclear morphology and lower LMNA/C expression. **A-B.** Representative confocal microscopy images of dHL60 cells stimulated with 100nM LTB_4_ for 15 min, fixed and immunostained for LBR (yellow) and LMNA/C (magenta) (**a)** and FLAP (green) (**b)**, both co-stained with DAPI (cyan). Scale bar is 10µm. **C.** Cartoon depicting the relationship between nuclear shape and morphological shape parameters on a scale from 0 to 1. Eccentricity increases and Form Factor decreases with nuclear lobulation. **D-E.** Dot plot graphs showing Form Factor (**d**) and Eccentricity (**e**) of nuclear shape. Data points represent pooled means per well of 24-well plate from three independent experiments. *P* values calculated using two-tailed Mann-Whitney test (**d**) or two-tailed t test (**e**). Data presented as mean ± s.e.m. **F-H.** Bar graphs showing normalized intensity of LBR (**a**), LMNA/C (**b**), and FLAP (**c**) in nuclei of dHL60 cells. Data points represent pooled means per well of 24-well plate from three independent experiments. *P* values calculated using two-tailed Mann-Whitney test (**a**) or two-tailed t test (**b, c**). Data presented as mean ± s.e.m. **I-K.** Dot plot graphs showing normalized intensity of LBR (**i**), LMNA/C (**j**), FLAP (**k**) for all cells across three independent experiments. Black bar represents mean ± interquartile range.

## DISCUSSION

The discovery that HL60 cells can be differentiated into neutrophil-like cells has established them as an essential model for neutrophil biology research. However, the current differentiation protocols of HL60 cells do not fully recapitulate the morphological and functional characteristics of human primary neutrophils isolated from peripheral blood. ATRA differentiation results in cells with lobulated nuclei and low levels of LMNA/C^45,47^, but the cells are spontaneously apoptotic^40,41^, have low expression of chemoattractant receptors such as FPR1, and poor chemotactic activity and ROS generation in response to fMLF^53,54^. On the other hand, DMSO dHL60 cells have robust oxidative burst and migratory responses to fMLF^55,38^ but do not harbor lobulated nuclei ^39^ and their nuclei express high levels of LMNA/C^56,57^. In this study we found that sequential treatment with a low concentration of ATRA followed by DMSO compensates for the deficiencies of the singular treatments. A1D4 dHL60 cells are not spontaneously apoptotic, express surface markers of mature granulocytes, including FPR1, and have robust chemotaxis and oxidative burst in response to fMLF. In addition, a high proportion of these cells have multilobulated nuclei with a concomitant reduction in the expression of LMNA/C, compared to DMSO dHL60 cells.

ATRA is commonly used in the clinic to treat patients with Acute Promyelocytic Leukemia, in combination with chemotherapy, to induce differentiation of leukemic cells to mature granulocytes which then die off^58^. As a result, the mechanisms of ATRA-mediated differentiation of HL60 cells are well understood. ATRA induces transcriptional changes through its interaction with the nuclear receptor heterodimer of RARα and retinoid x receptor (RXR). This alters the transcriptional landscape by displacing the RARα/RXR heterodimer from its basal repressive functions, allowing the transcription of genes essential for granulocytic differentiation, including the C/EBP family of transcription factors^59,60^. ATRA treatment has also been shown to block the transcription of c-Myc, which is typically overexpressed in HL60 cells and essential for their continued proliferation^4,61^. After extended treatment, ATRA can also activate phosphatidylinositol (PI) 3’-kinase^62^, *src*-family protein-tyrosine kinases^63^ and mitogen-activated protein (MAP) kinase signaling pathways^64^ that arrest growth and ultimately induce morphological and cytoskeletal changes. However, these signaling cascades appear at later timepoints after ATRA treatment^65^, whereas transcriptional changes occur in a matter of hours^59^. In contrast, the mechanisms driving DMSO differentiation are considerably less well defined. DMSO treatment accompanies a decrease in the fluidity of cell membranes – indicating dramatic changes in lipid composition^20^. DMSO also causes changes in chromatin structure^66^ and activation of protein kinase C^14^, both resulting in a reduction in c-Myc. Overall, these effects may result in differentiation not just by blocking proliferation but also by priming the cell for functional responses such as chemotaxis. We reason that sequentially treating HL60 cells for one day with ATRA and four days with DMSO induces the transcriptional effects of ATRA treatment at early time points while activating DMSO-mediated cell priming at later time points. This could explain how our method gives rise to dHL60 cells with a more mature granulocytic phenotype and robust immune functions.

Nuclear segmentation and high chemotactic motility are key identifiers of the neutrophil cell type^46,67^. In addition, interest has grown in recent years to study the role of the nucleus in migration and signaling^46,68^. As a widely adopted model for neutrophil studies, HL60 cells serve as an important model for understanding the molecular mechanisms of neutrophil function. In this context, our method improves the phenotypic and functional relevance of dHL60 cells as a model to study neutrophil biology.

## ACKNOWLEDGMENTS AND SOURCES OF FUNDING

We are thankful to members of the Parent laboratory for advice and support. We also thank Dr. Jonathan Sexton (University of Michigan) for assistance and use of the high-content confocal microscope. The funders had no role in study design, data collection and analysis, decision to publish or preparation of the manuscript. This work was supported by funding from the University of Michigan School of Medicine (CAP), predoctoral fellowship award from an American Heart Association (AWD025905) (SPC), and by several grant awards from National Institute of Health, namely: T32 training program in Cell and Molecular biology GM145470 (SPC) and R01AI152517 (CAP).

## AUTHORSHIP CONTRIBUTION STATEMENT

Conceptualization: SPC and CAP

Investigation: SPC, GY, CR and SSR

Data curation - formal analysis: SPC and SSR

Project Administration/oversight : CAP

Writing - original draft: SPC

Writing - review, editing, and revision: SPC, GY, CR, SSR, and CAP

## SUPPLEMENTARY INFORMATION

**Supplementary Figure 1.**
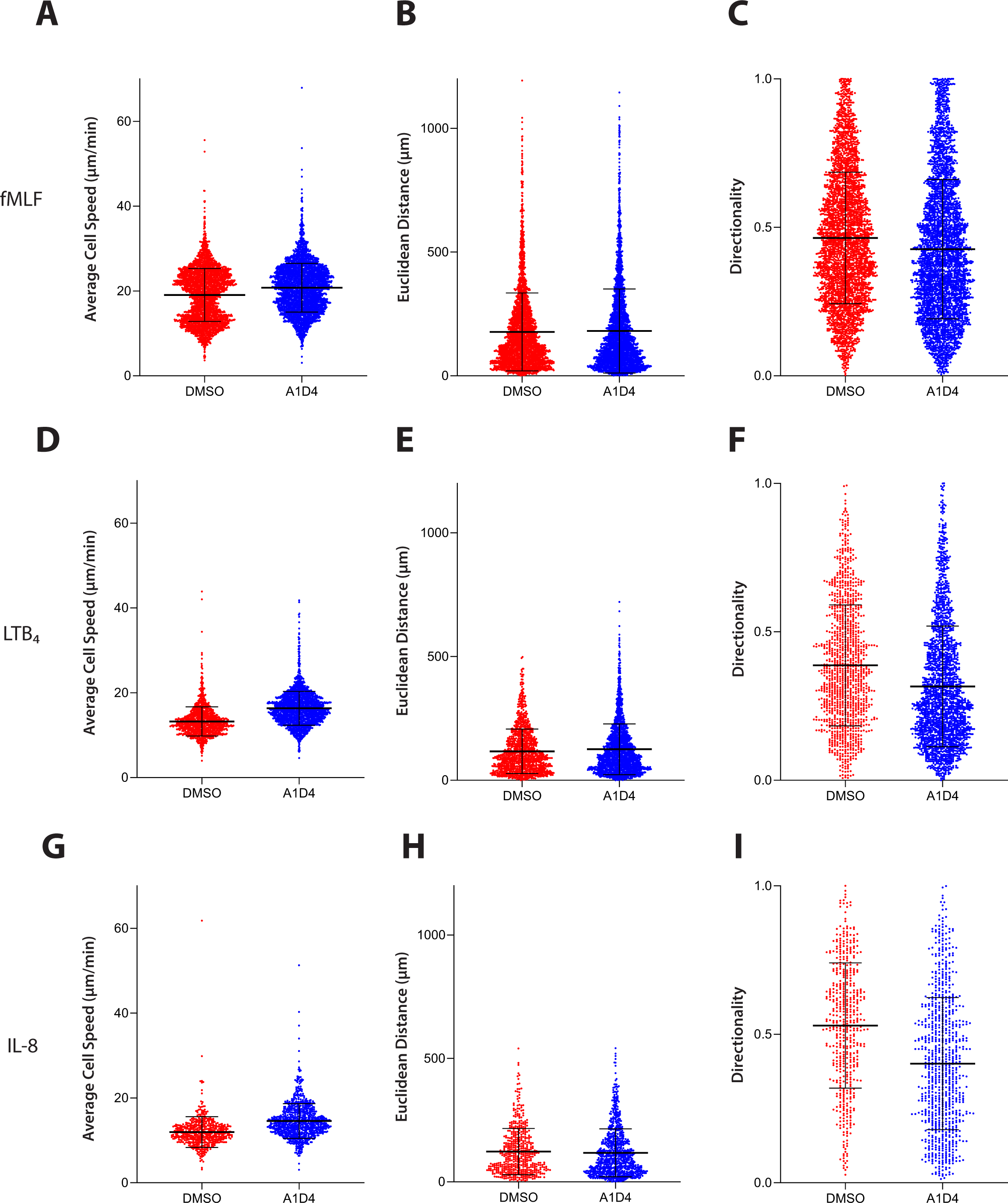
Chemotaxis parameters for individual DMSO and A1D4 dHL60 cells. **A-I.** Dot plot graphs showing chemotaxis parameters after 60 min of migration towards fMLF (**a-c**), LTB_4_ (**d-f**) N=3, IL-8 (**g-i**) N=3. For each chemoattractant the following parameters are displayed: average cell speed (**a, d, g**), which represents cells’ average speed during migration from each independent experiment; Euclidean distance travelled (**e, i, m**), which shows cells’ distance traveled in a straight line; and directionality (**f, j, n**), which represents the ratio of Euclidean distance to total accumulated distance traveled. Black bar represents mean ± interquartile range.

**Supplementary Figure 2.**
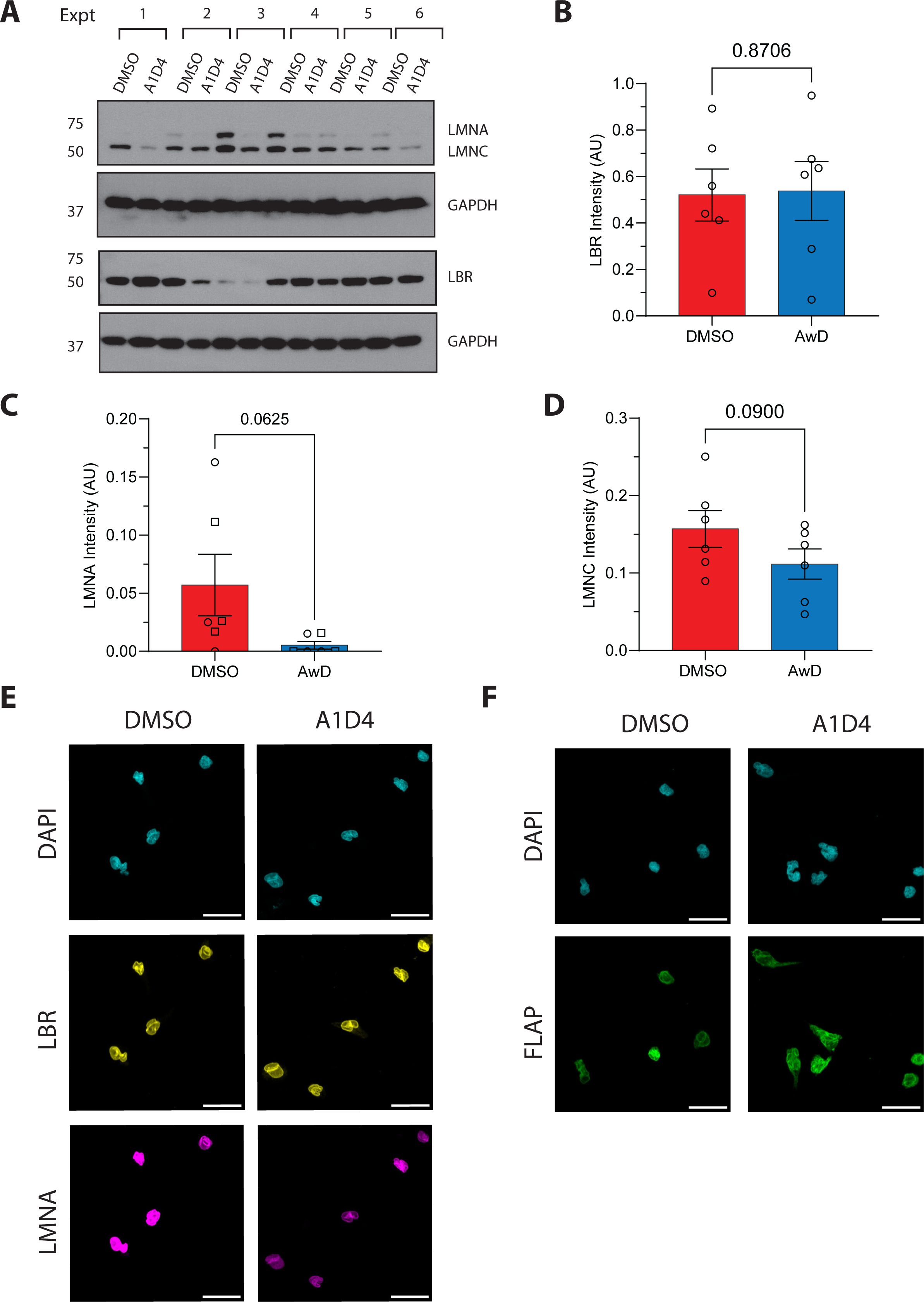
Nuclear protein expression in dHL60 cells. **A.** Western blot showing levels of LMNA, LMNC, LBR and GAPDH in cell lysates from DMSO and A1D4 dHL60 cells. Molecular weights (kDa) are indicated on the left. N=6. **B-D.** Bar graphs showing expression levels via band intensity of LBR (**b**), LMNA (**c**), and LMNC (**d**) normalized to GAPDH expression. *P* values calculated using two-tailed t test. Data are presented as mean ± s.e.m. **E-F.** Representative microscopy images of multiple fixed dHL60 cells stimulated with 100nM LTB_4_ for 15 min, immunostained with LBR (yellow) and LMNA/C (magenta) in **e** and FLAP (green) in **f**, both co-stained with DAPI (cyan). Scale bar is 20µm.

**Supplementary Video 1**

Representative widefield microscopy movies of dHL60 cells migrating for 60 min toward fMLF. Nuclei are visualized by a purple circle and cell tracks are shown as a trailing, colored line. The associated differentiation condition and time stamp are provided at the top left and right of each movie panel. Scale bar is 200µm. Representative of three independent experiments.

**Supplementary Video 2**

Representative widefield microscopy movies of dHL60 cells migrating for 60 min towards fMLF, LTB_4_, or IL-8. Nuclei are visualized by a purple circle and cell tracks are shown as a trailing, colored line. The associated differentiation condition is provided on the left of the two rows of movies. The chemoattractant and time stamp are provided in the bottom and top right of each movie panel. Scale bar is 200µm. Representative of three independent experiments.

